# Pachytene karyotypes of 17 species of birds

**DOI:** 10.1101/2022.01.01.473627

**Authors:** Anastasia Y. Slobodchikova, Lyubov P. Malinovskaya, Ekaterina O. Grishko, Inna E. Pristyazhnyuk, Anna A. Torgasheva, Pavel M. Borodin

## Abstract

Karyotypes of less than 10% of bird species are known. Using immunolocalization of the synaptonemal complex, the core structure of meiotic chromosomes at the pachytene stage, and centromere proteins we described male pachytene karyotypes of seventeen species of birds. This method enables higher resolution than the conventional analyses of metaphase chromosomes. We provided the first descriptions of the karyotypes of three species (Rook, Blyth’s reed warbler and European pied flycatcher), corrected the published data on the karyotypes of ten species and confirmed them for four species. All passerine species examined have highly conservative karyotypes, 2n=80-82 with seven pairs of macrochromosomes and 33-34 pairs of microchromosomes. In all of them but not in the Common cuckoo we revealed single copies of the germline restricted chromosomes varying in size and morphology even between closely related species. This indicates a fast evolution of this additional chromosome. The interspecies differences concern the sizes of the macrochromosomes, morphology of the microchromosomes and sizes of the centromeres. The pachytene cells of the Gouldian finch, Brambling and Common linnet contained heteromorphic synaptonemal complexes indicating heterozygosity for inversions or centromere shifts. The European pied flycatcher, Gouldian finch and Domestic canary have extended centromeres in several macro- and microchromosomes.

## Introduction

Birds provide an interesting model to study chromosome evolution. They have undergone rapid speciation and evolved various adaptations to a wide variety of habitats. Yet, their karyotypes (chromosome sets) are very conservative. They are bimodal, i.e. composed of macro- and microchromosomes, the feature inherited from their reptilian ancestors^1,2^. Diploid chromosome number (2n) of most karyotypes of bird species is about 78–82^3^. The number of chromosome arms, so-called fundamental number (FN), also vary in narrow limits: 90-110. Unfortunately, less than 10% of bird species have been karyotyped. In passerines, the most speciose and diverse order of birds, the portion of karyotyped species is even smaller – 7%^4^. Paradoxically, the karyotype is unknown even in the species whose genomes have been sequenced, annotated and studied in detail, such as the European pied flycatcher *Ficedula hypoleuca*^5^.

Most avian karyotypes have been described in the 1960-1970s with the use of conventional methods of chromosome preparation and staining available at that time^3^. Development of the methods of chromosome analysis at the pachytene stage of meiosis provided a rather efficient karyotyping tool^6^. These methods are based on visualization of the synaptonemal complex (SC), the structure that mediates synapsis and recombination of homologous chromosomes. SC is composed of two lateral elements, to which the chromatin loops are attached, and the central element that holds homologous chromosomes together. At the pachytene stage of meiotic prophase, the chromosomes are less compacted than at metaphase^7^. Therefore, analysis of SC provides higher resolution than the conventional analyses of metaphase chromosomes. This is especially important in the cytogenetics studies of the bimodal karyotypes because a morphology of the microchromosomes and even their number are rather difficult to assess at the conventional chromosome spreads^8,9^. The SC analysis is particularly efficient in the detection of heterozygosity for all types of chromosome rearrangements: inversions, translocations, deletions and duplications. The heterozygotes for the structural variants and heterogametic organisms produce characteristic heteromorphic SCs^10–12^.

Analysis of pachytene chromosomes led to the discovery of germline-restricted chromosome (GRC)^13^. The GRC was present in germline cells and absent in somatic cells in all 18 species of passerine birds examined so far^14,15^. It has not been found in any non-passerine bird^14^. In female germ cells, GRC is usually present in two copies, which synapse and recombine with each other. At the meiotic prophase, the GRC bivalents are practically indistinguishable from normal autosomal bivalents. They can only be revealed by a comparative subtractive analysis of pachytene and somatic karyotype. In male germ cells, GRC is usually present in one copy, which forms a univalent at pachytene, easily distinguishable from the bivalents of the autosomes and ZZ by its thin, coiled and often fragmented SC. The GRC univalents in male pachytene cells are always surrounded by dense chromatin clouds heavily labeled by anticentromere antibodies. Among the Passeriformes, the GRC varies in size and morphology^13,14,16,17^

In this study, using immunolocalization of SYCP3, main protein of the lateral elements of synaptonemal complex (SC) and centromere proteins we examined male pachytene karyotypes of sixteen passerine species with special attention to the GRC and one outgroup species the Common cuckoo *Cuculus canorus.* The sources of the specimens are shown in Supplementary Table 1. In each specimen we photographed, thoroughly examined and measured at least 20 well-spread pachytene nuclei containing complete chromosome sets. The measurements were used to generate idiograms. We compared the pachytene karyotypes with the karyotypes described earlier and mostly obtained by conventional methods of chromosome preparation and staining.

## Results

Figure 1 shows microphotographs of the SC spreads after immunolocalization of SYCP3 (red) and centromere proteins (blue) of all species examined. Figure 2 shows idiograms of pachytene karyotypes of the studied species. The haploid chromosome number is equal to the number of the SCs (n), the haploid number of the chromosome arms is equal to one for each acrocentric chromosome and to two for each meta- and submetacentric chromosome (Fn), the total SC length and brief description of GRC are shown in Table 1.

**Figure 1.**
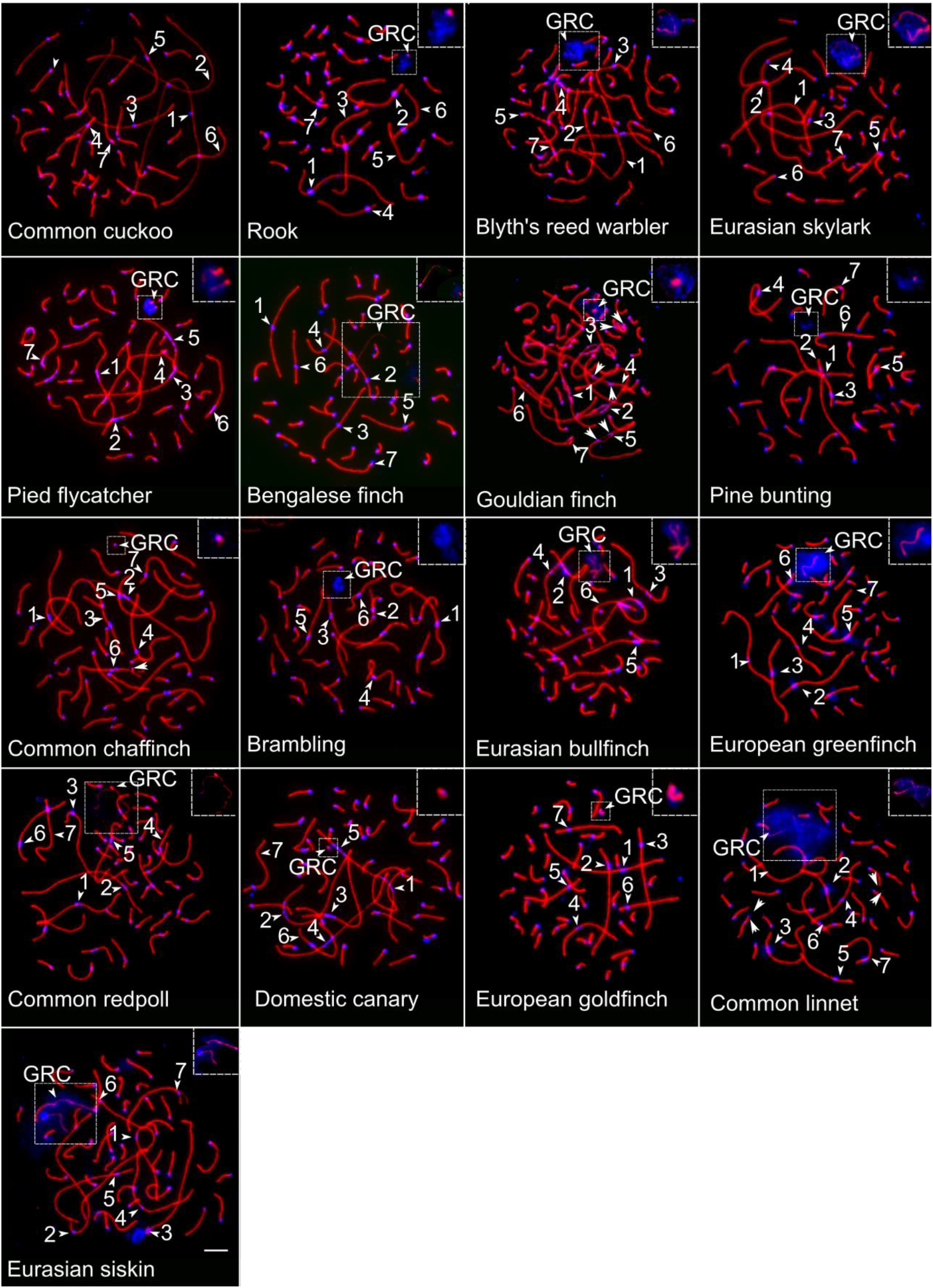
Microphotographs of the SC spreads after immunolocalization of SYCP3 (red) and centromere proteins (blue) of all species examined. The inserts show close-up of the GRC. Arrowheads indicate centromeres, arrows show heteromorphic bivalents. Numbers indicate SCs of the macrochromosomes. Bar—5 μm.

**Figure 2.**
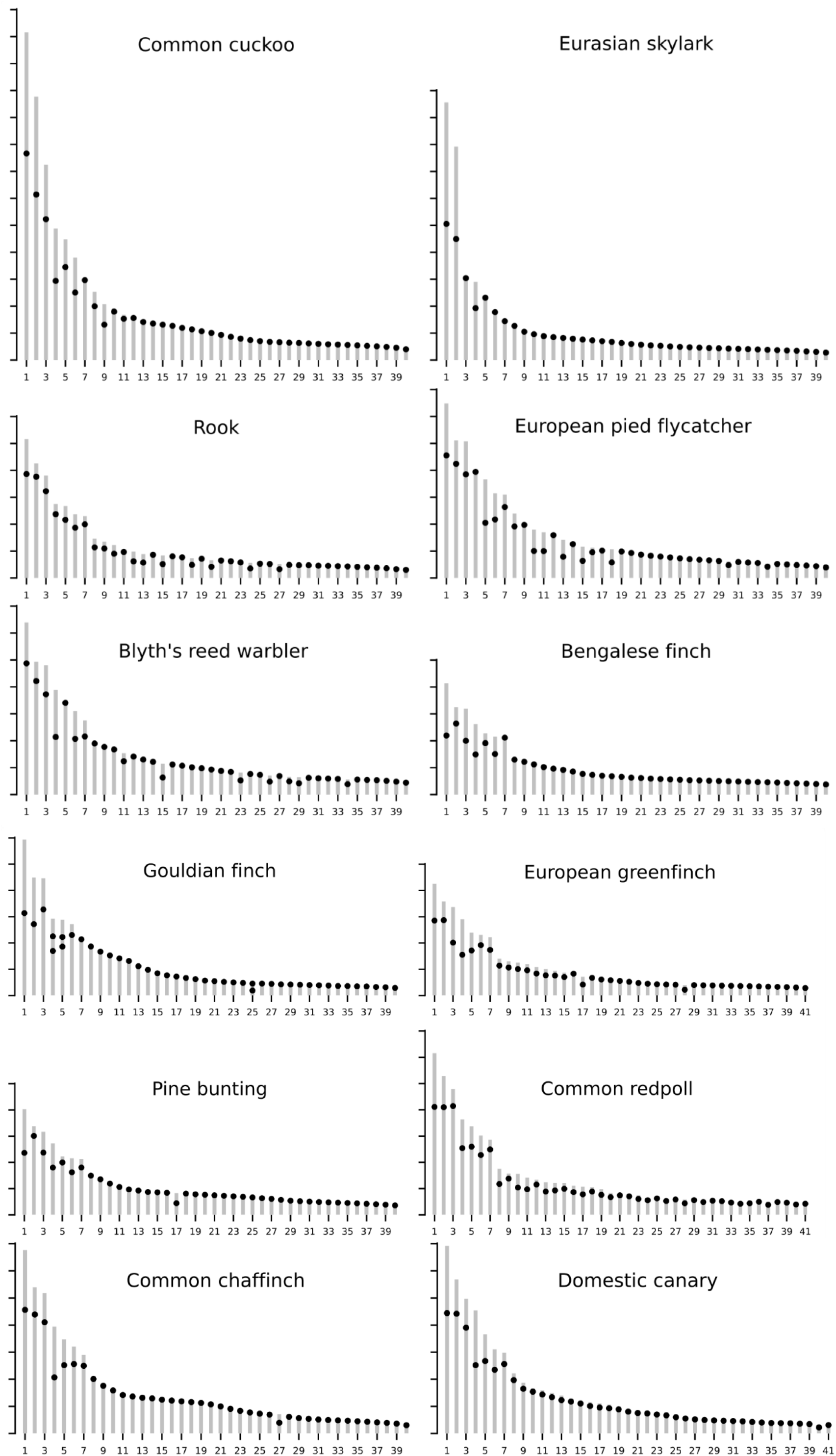
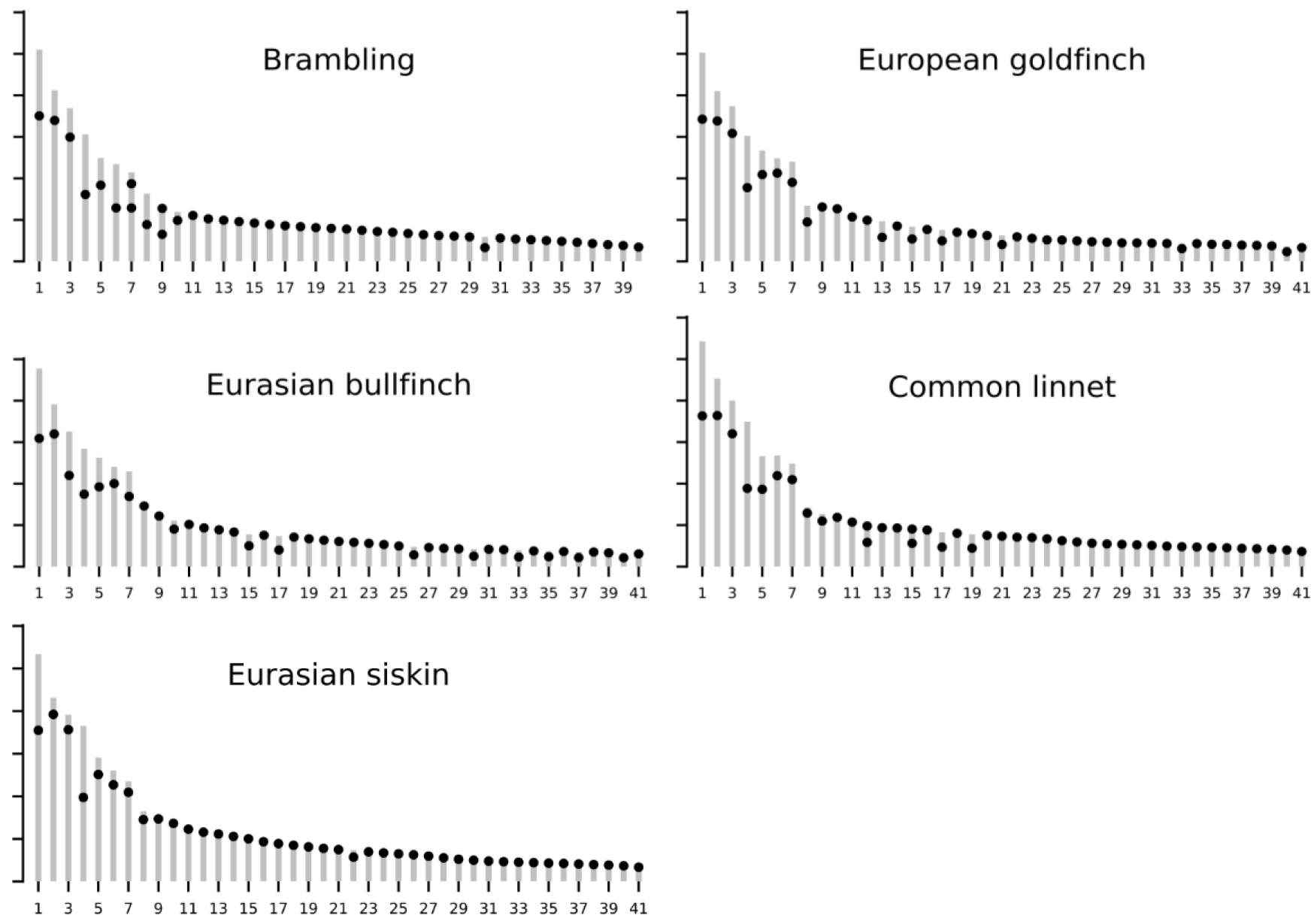
Idiograms of pachytene karyotypes without GRCs of the studied species. The *y*-axes show average SC length in μm. The x-axes SCs in decreasing size order. Black circles indicate centromeres. Arrows indicate additional centromeres in the heteromorphic bivalents

**Table 1.** Haploid chromosome numbers (n), haploid numbers of the chromosome arms (Fn), total SC length of the basic chromosome set and characteristics of GRC of 17 bird species

| Common name | n | Fn | SC length (μm) | No of cells analyzed | GRC |
| --- | --- | --- | --- | --- | --- |
| Common cuckoo | 40 | 48 | 387±68 | 60 | Absent |
| Rook | 40 | 57 | 226±29 | 60 | Micro |
| Blyth's reed warbler | 40 | 53 | 297±51 | 63 | Macro |
| Eurasian skylark | 40 | 43 | 241±75 | 56 | Macro |
| European pied flycatcher | 40 | 55 | 309±47 | 175 | Micro |
| Bengalese finch | 40 | 46 | 208±24 | 57 | Macro |
| Gouldian finch | 40 | 46 | 238±34 | 27 | Micro |
| Pine bunting | 40 | 48 | 216±33 | 60 | Micro |
| Common chaffinch | 40 | 48 | 301±38 | 36 | Micro |
| Brambling | 40 | 50 | 237±19 | 34 | Micro |
| Eurasian bullfinch | 41 | 57 | 209±47 | 78 | Macro |
| European greenfinch | 41 | 58 | 209±17 | 49 | Micro |
| Common redpoll | 41 | 70 | 287±31 | 31 | Macro |
| Domestic canary | 41 | 56 | 302±41 | 30 | Micro |
| European goldfinch | 41 | 55 | 220±29 | 23 | Micro |
| Common linnet | 41 | 53 | 246±29 | 37 | Macro |
| Eurasian siskin | 41 | 50 | 245±66 | 108 | Macro |

The descriptions of the chromosome morphologies are based on the estimates of their centromeric indices. Those with the centromeric index close to 0 is scored as the acrocentrics, between 0 and 0.4 as the submetacentrics, and between 0.4 and 0.5 as the metacentrics. In all species, we identified the seven largest SCs as macro-SC, the others – as micro-SCs.

We estimated the size of GRC based on the size of GRC chromatin labeled by anticentromere antibodies and the length of the lateral element of GRC SC if it was completely formed. We classified GRCs of size comparable to macrochromosomes of the basic set as macro-GRCs, others – as micro-GRCs. In most species, we found a variation in the SC appearance of macro- and micro-GRCs between the cells. It could form a complete, or fragmented, or dot-like lateral element of SC or do not form it at all.

### Common cuckoo *Cuculus canorus*

The somatic karyotype of the Common cuckoo has been described by Bian and Li^18^. They indicated 2n=78. The pachytene karyotype of this species comprises 40 chromosome pairs (2n = 80). The total length of its SC set is one of the largest among the bird species examined so far. The excess of the SC length is mainly due to very large metacentric SCs 1 and 2 and submetacentric SC3. Macro-SC4 is a metacentric. Macro-SCs 5 and 6 and micro-SCs 8 and 9 are submetacentrics, Macro-SC7 and all other 30 micro-SCs are acrocentrics. The Common cuckoo as well as all other non-passerine birds studied so far does not have GRC in its pachytene karyotype.

### Rook *Corvus frugilegus*

The somatic karyotype of the Rook has not been studied yet. The diploid number of the closely related species *Corvus corax* is 80^19^. Our results indicate that the basic diploid chromosome set of the Rook is 80. The pachytene karyotype of Rook comprises 40 bivalents and one univalent of a GRC surrounded by a small cloud of anticentromere antibodies. All of Rook’s macro-SCs are submetacentrics. Among the micro-SCs, there are ten submetacentrics and twenty-three acrocentrics. In most pachytene cells, GRC forms dot-like SC while in some cells SC is not formed.

### Blyth’s reed warbler *Acrocephalus dumetorum*

The somatic karyotype of the Blyth’s reed warbler remains unknown. The diploid number of the closely related species *Acrocephalus agricola* is 78^20^. Pachytene karyotype of the Blyth’s reed warbler comprises 40 chromosome pairs (2n = 80). Its macro-SCs 1, 2, 3, 6 and 7 are submetacentrics, SC4 is metacentric, SC5 is acrocentric. Most micro-SCs are acrocentrics. There are one metacentric and six submetacentric micro-SCs.

GRC in the Blyth’s reed warbler occurs as a large acrocentric univalent with fragmented SC surrounded by a chromatin cloud labeled by anticentromere antibodies. The level of fragmentation varies between the cells from evenly labeled SC to a dispersed series of short fragments.

### Eurasian skylark *Alauda arvensis*

The first description of the Eurasian skylark somatic karyotype was provided by Udagawa^21^ on the basis of histological sections of embryonic testes and ovaries. The spermatogonial 2n was estimated as 78, oogonial as 77. Li and Bian^22^ estimated 2n the Oriental skylark *Alauda gulgula* as 76±. We found that the pachytene karyotype of the Eurasian skylark contains 40 chromosome pairs (2n =80). In terms of SC sizes, it is extremely asymmetric. Metacentric SCs 1 and 2 comprise 36% of the total SC length. This is in agreement with the description of the Alaudidae karyotype by Li and Bian^22^. They described the Z- and W-chromosomes as the largest elements, metacentric and submetacentric respectively. The 1^st^ macrochromosome was the second largest metacentric.

One of the two largest SCs in the pachytene karyotype of the Eurasian skylark is probably the neo-Z chromosome. Genomic analysis indicates that Z chromosome of several skylark species resulted from fusions between parts of the chromosomes Z, 3, 4A and 5. It is the largest sex chromosome found in birds (about 200 Mb)^23,24^. Another exceptionally large chromosome has probably also evolved via several chromosome fusions. Surprisingly, the chromosome number of the Eurasian skylark is the same as in most songbirds. This means that the fusions leading to the formation of the two largest macrochromosomes had been preceded or followed by a series of chromosome fissions. Besides the two largest macrochromosomes, the karyotype of the Eurasian skylark contains one submetacentric macrochromosome. All other macrochromosomes and microchromosomes are acrocentrics.

GRC of the Eurasian skylark occurs as a large chromatin cloud heavily labeled with anticentromere antibodies. It forms thin, pale and fragmented SC.

### European pied flycatcher *Ficedula hypoleuca*

The genome of the European pied flycatcher has been sequenced, annotated and studied in detail^5^, yet its karyotype remains unknown. Its close relative *Ficedula parva* contains 40 chromosome pairs (2n=80)^25^. The pachytene karyotype of the European pied flycatcher contains 40 bivalents (2n=80). Almost all its macro-SCs are submetacentrics, SC4 is acrocentric, SC5 is metacentric. We are sure that SC4 is ZZ because analysis of pachytene oocytes of this species revealed heteromorphic ZW bivalent with acrocentric Z axis^26^. Among micro-SCs there are five metacentrics and four submetacentrics.

We observed extended pericentromeric regions in all meta- and submetacentric macro-SCs (but not in the acrocentric ZZ) and in several largest micro-SCs. They appeared as beads of several dots, which cover from 9% of SC1 to 41% of SC11/13 (Supplementary Table 2). The antibodies against the histone H3, di- and trimethylated at lysine 9 (H3K9me2/3), marking transcriptionally inactive chromatin, produced strong signals at the standard centromeres of the pachytene chromosomes, but not at the extended ones (Fig. 3a). In the corresponding somatic metaphase chromosomes of this species, we observed extended primary constrictions (Fig 3b).

**Figure 3.**
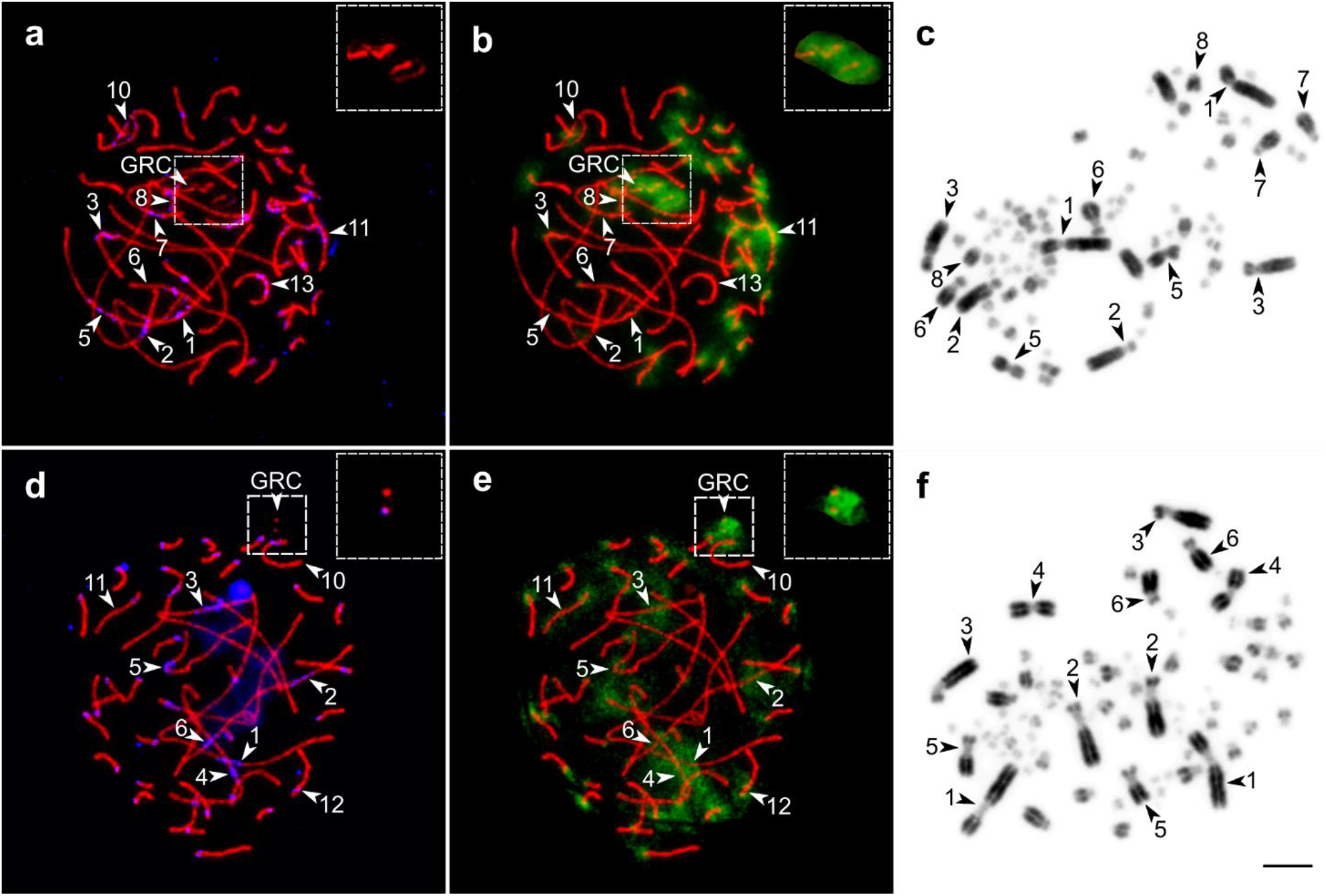
Pachytene spermatocytes immunolabeled with antibodies against SYCP3 (red) and centromere proteins (blue) (a,d) and H3K9me2/3 (green) (b, e); and somatic metaphase chromosomes stained with DAPI (c,f) of the European pied flycatcher (top row) and Domestic canary males (bottom row). The arrowheads indicate the extended centromeres and GRCs. The inserts show close-up of the GRCs. Bar—5 μm.

GRC in the European pied flycatcher usually appears as a chromatin cloud heavily labeled with anticentromere and H3K9me2/3 antibodies containing several SC fragments. All cells of one specimen contained one GRC. Another specimen was mosaic: 93 % of its pachytenes (126 out of 135) contained two univalents of GRC, the remaining cells contained one GRC. Earlier polymorphism and mosaicism for GRC number has been detected in the pale martin, great tit^16,17^.

### Bengalese finch *Lonchura striata domestica*

Pachytene karyotype of the Bengalese finch comprises 40 chromosome pairs (2n=80). This coincides with the description given by del Priore and Pigozzi^27^ and differs from the earlier description of its somatic karyotype as 2n =78^28–30^. Its SCs 1 and 4 are metacentrics, SCs 2, 3, 5 and 6 are submetacentrics. All other SCs are acrocentrics.

The Bengalese finch GRC has been first described by del Priore and Pigozzi^27^ and then by Torgasheva *et al.*^14^. It appears as the largest acrocentric univalent in the pachytene nucleus. We found a variation in the appearance of GRC SC between the cells: from complete lateral element to dot-like SC. Despite variation in the degree of SC polymerization, the cloud of anticentromere antibodies labeled GRC was similarly large in all cells, which allowed us to classify GRC of Bengalese finch as macro-GRC in accordance with the previous descriptions^14,27^

### Gouldian finch *Erythrura gouldiae*

Christidis^28^ described 2n of the Gouldian finch as 78. We found 40 chromosome pairs (2n=80) in pachytene cells of this species. SCs 1 and 2 are metacentrics, SCs 3 to 6 are submetacentrics. Macro-SC7 and all micro-SCs are acrocentrics. SCs 1, 2 and 3 have extended centromeres, similar to those described in the European pied flycatcher. They occupy around 13-20% of SC length (Supplementary Table 2). In half of cells analyzed, these regions are asynapsed whereas all other SCs are completely synapsed (Fig. 1). The delayed synapsis could be caused by an unusually extended pericentromeric heterochromatin.

We found heteromorphic SCs 4, 5 and 25. SC4 has two centromeres in metacentric and submetacentric positions. SC5 has two centromeres in different submetacentric positions. SC25 has centromeres in acrocentric and metacentric positions. GRC appears as a moderately sized chromatin cloud comparable to the largest microchromosomes labeled with anticentromere antibodies surrounding from one to three SC fragments.

### Pine bunting *Emberiza leucocephalos*

In the Pine bunting pachytene spermatocytes, we observed 40 bivalents (2n=80) while Radzhabli et al.^31^ and Bulatova^32^ described the diploid number of the somatic karyotype of this species as 2n=78. The pachytene karyotype of this species is rather similar to that of the Bengalese finch described above. There are three differences. SC4 is submetacentric, SC7 is submetacentric and one of the micro-SCs is metacentric. GRC of the Pine bunting is labeled by a small cloud of anticentromere antibodies. In most cells, GRC does not form the lateral element of SC, in some cells we detected a dot-like signal of anti-SYCP3 antibodies.

### Common chaffinch *Fringilla coelebs*

The Common chaffinch is an important model for evolutionary genetic studies. Its high-quality genome assembly has recently been published^33^. Our estimate of the Common chaffinch karyotype coincides with the earlier published one (2n= 80)^34^. Its pachytene karyotype comprises 40 chromosome pairs. Total SC length and the lengths of the macro-SCs in the Common chaffinch are much longer than in the closely related Pine bunting (Table 1). However, the morphology of their chromosomes is rather similar except SC1, which is submetacentric in the Common chaffinch.

GRC of the Common chaffinch forms dot-like SC labeled by a small cloud of anticentromere and H3K9me2/3 antibodies. To our surprise, in half of pachytene spermatocytes (26 out of 49 examined), we observed a univalent of a small microchromosome. Unlike the GRC, it was not labeled either with anticentromere or with H3K9me2/3 antibodies (Fig. 4a). We suggest that this univalent was originated by a premeiotic nondisjunction of one of the microchromosomes.

**Figure 4.**
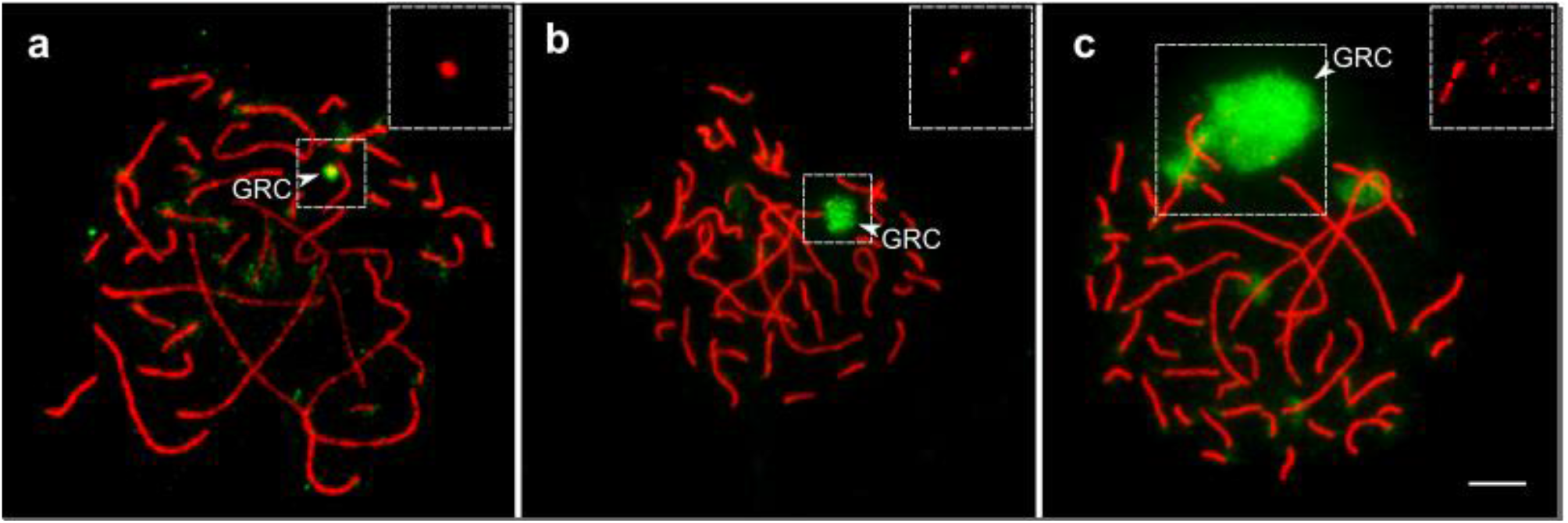
Pachytene spermatocytes of the Common chaffinch (a), Brambling (b) and Common linnet (c) after immunostaining with antibodies against SYCP3 (red) and H3K9me2/3 (green). The arrowheads indicate GRCs. The inserts show close-up of the GRCs. Bar—5 μm.

### Brambling *Fringilla montifringilla*

The somatic karyotype of the Brambling has been described as 2n=78^35^. We observed 40 pairs of chromosomes (2n=80) which were morphologically similar to the pairs detected in the Common chaffinch. There are a few differences. SC6 is metacentric, and two of micro-SCs are metacentric and one – is a submetacentric.

The specimen examined here has heteromorphic SCs 7 and 9. Each of them contains two centromeres in all cells examined. One homolog of chromosome 7 is the submetacentric as in most fringillids examined here. The other homolog of chromosome 7 is metacentric, which had probably resulted from pericentric inversion, as well as the metacentric homolog of the acrocentric chromosome 9 typical for the fringillids.

Similar to the GRC of the Pine bunting, the GRC of the Brambling is labeled by a small cloud of anticentromere antibodies and does not form the lateral element of SC in most analyzed cells. In some cells, we detected dot-like signals of anti-SYCP3 antibodies.

### Eurasian bullfinch *Pyrrhula pyrrhula*

Li and Bian^35^ described the somatic karyotype of the Eurasian bullfinch as 2n=78. The pachytene cells of this species contain 41 chromosome pairs (2n=82). All macro-SCs are submetacentric. Among micro-SCs we observed four metacentric and five submetacentric micro-SCs.

GRC of this species has been described by Torgasheva *et al.*^14^. It forms long acrocentric univalent evenly labeled by SYCP3 antibodies.

### European greenfinch *Chloris chloris*

The diploid chromosome number of the European greenfinch is 76 according to Hammar and Herlin^36^ and 80 according to Christidis^37^. We detected 41 bivalents (2n=82) in pachytene cells. The morphology of the seven largest macro-SCs is very similar to that described for the Eurasian bullfinch except for the morphology of macro-SCs 3 and 4. They are metacentric in the European greenfinch. The three smallest macro-SCs and eight largest micro-SCs are submetacentrics. There are also two metacentric micro-SCs while all the others are acrocentrics.

GRC of the European greenfinch appears as a partially formed lateral element of SC comparable in size to the largest microchromosomes and labeled by anticentromere antibodies.

### Common redpoll *Acanthis flammea*

According to Li and Bian^35^ diploid chromosome number of the Common redpoll is 78. According to our estimate, it is 82. All macro-SCs are submetacentric. Among micro-SCs, twenty-two are submetacentrics and twelve are acrocentrics.

The pachytene karyotype of the Common redpoll also contains large acrocentric univalent of GRC evenly labeled by antibodies to SYCP3. In some cells, the SC of GRC was fragmented.

### Domestic canary *Serinus canaria* forma *domestica*

The karyotype of the Domestic canary has been first described by Ohno *et al.*^38^ as 2n=80± and recently by Da Silva Dos Santos *et al.*^39^ and Kiazim *et al.*^40^ as 80. The Domestic canary genome has been assembled and annotated but not yet to a chromosome level^41^.

Pachytene cells of the Domestic canary contain 41 chromosome pairs (2n=82). Its macro-SCs are larger than those in the fringillids described above, although their morphology is similar. Macro-SC4 is metacentric, all other macro-SCs are submetacentric. Eight micro-SCs are submetacentrics. All other micro-SCs of the Domestic canary are acrocentrics.

The Domestic canary shows unusually extended centromeres of macro-SCs and the largest micro-SCs, similar to that detected in the European pied flycatcher and the Gouldian finch. They were about 2-3 μm long covering about 10% of the SCs 1, 2/3, 4, 6, and about 15% of the SCs 5 and 10/11/12 (Fig. 3c, Supplementary Table 2). The antibodies against H3K9me2/3 label both the standard and extended centromeres of the pachytene chromosomes (Fig. 3c). We also detected substantially extended primary constrictions at the corresponding metaphase chromosomes (Fig. 3d).

GRC of this species has been described by Torgasheva et al.^14^. It appears as a small cloud of anticentromere antibodies with the dot-like lateral element of SC.

### European goldfinch *Carduelis carduelis*

Christidis^37^ described the somatic karyotype of the European goldfinch as 2n=82. Our data confirm this diploid chromosome number. We observed 41 bivalents in the pachytene cells of the European goldfinch. Its seven largest macro-SCs were smaller than those of the Domestic canary but similar in morphology. We observed one metacentric and six submetacentric micro-SCs, all other micro-SCs are acrocentric.

GRC of the European goldfinch has been described by Torgasheva et al.^14^ as micro-GRC. In most cells, GRC was present as one or two dot-like lateral SC fragments labeled by a small cloud of anticentromere antibodies. In some cells, GRC formed a complete lateral SC element.

### Common linnet *Carduelis cannabina*

Bulatova^20,32^ found 82 chromosomes in bone marrow cells of the Common linnet. This is in good agreement with our observation. We found 41 bivalents. Pachytene karyotypes of the Common linnet and the European goldfinch are almost the same. The difference concerns the morphology of the micro-SCs. The Common linnet has only two homomorphic metacentric and two heteromorphic metacentric/acrocentric micro-SCs. Micro-SC9 is submetacentric. All other micro-SCs are acrocentrics.

The heteromorphic SCs 12 and 15 contain two centromeres in all cells examined: one in acrocentric and one in metacentric positions. The metacentric homologs had probably resulted from pericentric inversions.

GRC of the Common linnet occurs as an acrocentric univalent with fragmented SC labeled by a large cloud of anticentromere antibodies.

### Eurasian siskin *Spinus spinus*

The diploid chromosome number in the Eurasian siskin was estimated as 78^30,32,35,42^. We found 41 chromosome pairs in the pachytene cells of this species (2n=82). The morphology of the macro-SCs is rather similar to that in the two fringellid species described above. However, the karyotype of the Eurasian siskin contains only two submetacentric micro-SC, all other micro-SCs are acrocentrics.

Torgasheva et al.^14^ classified its GRC as macro-GRC. In our spreads, it occurs as a large cloud of anticentromere antibodies with fragmented SC.

## Discussion

The main results of this study are:

1. the first description of the karyotypes of three species (the Rook, Blyth’s reed warbler and European pied flycatcher),
2. the correction of the published data on the karyotypes of ten songbird species,
3. the first detection of the extended centromeres in three model species (the European pied flycatcher, Gouldian finch and Domestic canary),
4. the first detection of chromosomal polymorphism in three species (the Gouldian finch, Brambling and Common linnet),
5. and detailed characterization of the GRCs of all songbird species examined

Why are we sure that our descriptions of the karyotypes are more precise and reliable than the published data based on conventionally prepared and stained somatic metaphase spreads?

There are at least four reasons for this. First, the pachytene chromosomes of birds are about 2-3 times longer than somatic metaphase chromosomes (see Fig. 3). Second, each object, which we identified at the microphotography as the bivalent of the basic chromosome set, was simultaneously labeled with two different antibodies: against SYCP3, the main protein of the lateral elements of the SC, and the centromere proteins. Univalents were distinguished from the bivalents by less intense SYCP3 labeling. Therefore a misidentification of the microchromosome as cell debris and the cell debris as a microchromosome was unlikely. Third, these three criteria were applied to at least 20 well-spread pachytene cells containing complete chromosome sets. This made our estimates statistically sound. Forth, the bivalents were not just counted, they were measured. This made possible an objective estimate of the morphology of the macrochromosomes and, what is more important and almost impossible at the somatic metaphases, the morphology of the microchromosomes.

A comparison between our and previously published chromosome numbers of the examined species shows that the earlier researchers always undercounted one or two chromosome pairs (Table 1). These errors were due to the very small sizes of the smallest microchromosomes and the relatively low specificity of the chromosome dyes. For these reasons, it was almost impossible to estimate the morphology of the small microchromosomes at the mitotic metaphase spreads. It was believed that of all them were acrocentric^42^. Our data show that this is true for the Common cuckoo, Eurasian skylark and Bengalese finch. Other songbirds have at least one meta- or submetacentric microchromosome, and most of the species have many of them.

Using anticentromere antibodies, we revealed unusual extended centromeres in almost all macrochromosomes and the largest microchromosomes of three model species (the European pied flycatcher, Gouldian finch and Domestic canary). They are visible both at the pachytene and mitotic metaphase spreads of these species. They are also visible in the published images of the Domestic canary metaphase chromosomes and shown at the idiograms^39,40^ but did not attract much attention.

Such long centromeres are rare. They have been detected in a few species of legumes^43,44^, fire ants^45^, Indian muntjac^46^ and marsupial hybrids^47^. However, they have not yet been described in birds. Robertsonian translocations and centromere drive have been suggested among the possible causes of the centromere extension^45,46^. The results of H3K9me2/3 immunolocalization indicate a variation in the epigenetic status of the extended centromeres. In the Domestic canary, both extended and standard centromeres are H3K9me2/3-positive indicating their heterochromatic state. In the European pied flycatcher, the standard centromeres are H3K9me2/3-positive, while the extended centromeres are H3K9me2/3-negative. Genetic composition or the extended centromeres deserve special attention.

One more advantage of the SC analysis is its high efficiency in the detection of the structural heterozygosity. In this study, we detected heteromorphic SC in the Gouldian finch, Brambling and Common linnet. In all cases, the bivalents displayed two misaligned centromeres. This may indicate heterozygosity either for pericentric inversions or for centromere repositions (centromere shift)^48^. Both types of chromosome rearrangements are implicated for karyotypic macroevolution of birds^40,49^. The inversions play an especially important role in the restriction of the gene flow between sympatric and parapatric species^50^. However, the intraspecific polymorphism for inversion is poorly studied in birds. Our findings indicate the targets for future studies.

Our study revealed a wide variation in GRC size and appearance. Torgasheva et al.^14^ suggested to classify them as macro- and micro-GRCs to fit the criteria for macro- and microchromosomes of the basic set. Indeed, all analyzed chromosomes fell into one of these categories. However, some micro-GRCs were much smaller than the smallest microchromosomes of the basic set that can be inferred from the size of the chromatin cloud labeled by anticentromere and H3K9me2/3 antibodies. This variation and the lack of phylogenetic clustering by size confirm the highly dynamic nature of GRC^14^. It was shown that GRCs of different species contain different multiply repeated sequences, which probably can be accumulated and/or be lost rather quickly^51^.

The SC of GRCs in pachytene spermatocytes of most species examined here appeared fragmented, however, the degree of fragmentation varied between cells. It is unclear if this feature is related to the different properties of GRCs in different species (its genetic content, the degree of heterochromatinization) or the intercellular and interindividual variation in the effectiveness of cohesin loading and SC polymerization since for most birds studied to date, only one sample was analyzed. The interindividual variation in GRC appearance was observed in pachytene spermatocytes of the Great tit^17^. The intense chromatin labeling with antibodies to centromere proteins and H3K9-modified histone allowed us to distinguish between GRC and accidental autosomal univalents.

The results of our study indicate several lines of future research. To minimize sacrificing birds we described the karyotype of most species by single specimen examined. The sample size might be increased at least for the most interesting species by targeted examination of the somatic karyotypes, which can be obtained from short-term fibroblast cultures derived from blood or feather pulp.

It seems important to estimate the frequency, geographic distribution and probable adaptive significance of the inversion polymorphism detected in the Gouldian finch, Brambling and Common linnet, the species with wide breeding and residence areas.

Another interesting species is the Eurasian skylark with two giant chromosomes. The origin of its Z/W chromosomes has been resolved^23,24^. The genetic content of another giant chromosome of the Eurasian skylark remains unknown. FISH with universal BAC-probes^52^ might shed a light on its origin. The microchromosomes of this species also deserve close attention because despite the fusions of several macrochromosomes in the neo-Z and in another giant chromosome its chromosome number remains the same as in most songbirds.

The nature, evolution and adaptive significance of the extended centromeres of Gouldian finch, European pied flycatcher, and Domestic canary are of special interest. Recent advances in sequencing and bioinformatic analysis of the repetitive DNA of birds^53^ make it possible to address these questions.

## Methods

### Specimens

Adult males were sampled at the beginning of the breeding season (April-May). The sources of the material, the number of specimens are shown in Supplementary Table 1. The birds were handled and euthanized in accordance with the approved national guidelines for the care and use of laboratory animals.

### Spermatocyte spreading and immunostaining

Chromosome spreads were prepared by the drying down method^54^. Testes were dissected and placed in an extraction buffer for 30-60 min. Small pieces of testis were macerated in 40 μl of 0.1M sucrose on a clean glass slide. A drop of the fine suspension was dropped at the slide dipped in 1% paraformaldehyde solution (Sigma-Aldrich, cat# 158127), pH 9.2. The slides were incubated for 2 h in a humid chamber, washed in 0.4% Kodak Photo-Flo 200 (Kodak, cat# 742057), dried at room temperature and kept in sealed containers at −20°C until use.

Immunostaining was performed according to Anderson et al.^55^ using rabbit polyclonal anti-SYCP3 (1:500; Abcam, cat# ab15093), human anticentromere (1:100; Antibodies Inc., cat# 15-234) and mouse monoclonal anti-H3K9me2/3 (1:100, Cell Signaling, cat# 5327) primary antibodies. The secondary antibodies used were Cy3-conjugated goat anti-rabbit (1:500; Jackson ImmunoResearch, cat# 111-165-144), AMCA-conjugated donkey anti-human (1:100; Jackson ImmunoResearch, cat# 709-155-149) and FITC-conjugated goat anti-mouse (1:100; Jackson ImmunoResearch, cat# 115-095-003). The slides were incubated overnight with primary antibodies and 1 h with secondary antibodies at 37°C in a humid chamber. Slides were mounted in Vectashield antifade mounting medium (Vector Laboratories, cat# H-1000-10, United States).

### Mitotic metaphase chromosome preparations

The mitotic metaphase chromosomes were obtained from the cell culture of the gonads. The primary cell cultures were established from the minced gonads, that were successively treated with 100ng/ml Collagenase I (Sigma, cat# SCR103) and 0.25% Trypsin-EDTA (Sigma, cat# T4174) for 20 min by every component at 37 °C. The cell cultures were maintained in Dulbecco’s Modified Eagle’s medium (Gibco, cat# 41965039) supplemented with 10% Fetal bovine serum (Gibco, cat# 10270106), 2% Chicken serum (Gibco, cat# 16110082), GlutaMAX supplement (TermoFisher, cat# 35050038) and penicillin/streptomycin (Sigma-Aldrich, cat# P0781). Preparation of metaphase chromosomes from the bird’s cell culture was performed at the 2d to 3d passage according to modified protocol of chicken spermatogonial and follicles cell culture^56,57^. Briefly, the cell culture was incubated in a culture medium with the addition of 0.1 μg/ml colchicine for 3 h and collected by trypsin treatment. The cells were treated with hypotonic solution (0.56% KCl) for 30 min and fixed in 3:1 Methanol: Glacial Acetic Acid. Slides were mounted in Vectashield antifade mounting medium with DAPI (Vector Laboratories, cat# H-1200, United States).

### Microscopic image analysis

The preparations were visualized with an Axioplan 2 imaging microscope (Carl Zeiss) equipped with a CCD camera (CV M300, JAI), CHROMA filter sets, and the ISIS4 image-processing package (MetaSystems GmbH). In each specimen at least 20 well-spread pachytene nuclei containing complete chromosome sets were photographed. The brightness and contrast of all images were enhanced using Corel PaintShop Photo Pro X6 (Corel Corp).

### Chromosome measurements and generation of idiograms

The centromeres were identified by anticentromere antibodies. The length of the SC of each chromosome arm was measured in micrometers and the positions of centromeres were recorded using MicroMeasure 3.3^58^. We plotted idiograms after measuring at least 30 cells for each species. In each cell, we sorted SCs by their length in μm, identified those that can be distinguished unambiguously by the length and centromere index (most macrochromosomes in most species) and measured the average of these parameters across all cells. We grouped SCs that could not be unambiguously distinguished by the length and/or centromere indices (most microchromosomes in most species), ranged them within the group and measured the average parameters for each rank. Statistica 6.0 software package (StatSoft) was used for descriptive statistics.

## Declarations

### Ethics approval

The birds were handled and euthanized in accordance with the approved national guidelines for the care and use of laboratory animals. All experiments were reviewed and approved by the Animal Care and Use Committee of the Institute of Cytology and Genetics SB RAS (protocol # 114 of 17 December 2021).

## Acknowledgements

We thank the Core Facility for Microscopy of Biologic Objects, SB RAS for granting access to microscopic equipment.

## Funding

The study and open access fee were funded by Russian Science Foundation, grant number 20-64-46021. Microscopy was carried out at the Core Facility for Microscopy of Biologic Objects, SB RAS, Novosibirsk, Russia (regulation no. 3054). The funding bodies play no role in the design of the study and collection, analysis, and interpretation of data and in writing the manuscript.

## Authors’ contributions

A.T. and P.B. provided the research idea and designed the experiments. L.M., A.S., E.G., and I.P. conducted the experiments, collected the data, finished the data analysis and compiled the results. A.T. and P.B. supervised the research and wrote the main manuscript text. All authors reviewed and approved the final manuscript.

## Competing interests

The authors declare that they have no competing interests.

## Availability of data and materials

Publicly available datasets were analyzed in this study. These data can be found here: https://meiosislab.com/projects/chromosomes/17caryo.xls.

## Supplementary materials

Supplementary Table 1. Sources of the birds examined

Supplementary Table 2 Size of expanded centromeres

## Supplementary materials

**Supplementary Table 1.** Sources of the birds examined.

| Common name | No of specimens | Source |
| --- | --- | --- |
| Common cuckoo | 1 | Provided by Bird Rehabilitation Centre after fatal accident trauma |
| Rook | 1 | Provided by Bird Rehabilitation Centre after fatal accident trauma |
| Blyth's reed warbler | 1 | Trapped in Novosibirsk |
| Eurasian skylark | 1 | Trapped in Novosibirsk |
| European pied flycatcher | 2 | Trapped in Novosibirsk |
| Bengalese finch | 1 | Purchased a commercial breeder |
| Gouldian finch | 1 | Purchased a commercial breeder |
| Pine bunting | 1 | Trapped in Novosibirsk |
| Common chaffinch | 1 | Purchased from a commercial breeder |
| Brambling | 1 | Purchased from a commercial breeder |
| Eurasian bullfinch | 2 | Provided by Bird Rehabilitation Centre after fatal accident trauma |
| European greenfinch | 1 | Purchased from a commercial breeder |
| Common redpoll | 1 | Purchased from a commercial breeder |
| Domestic canary | 1 | Purchased from a commercial breeder |
| European goldfinch | 1 | Purchased from a commercial breeder |
| Common linnet | 1 | Purchased from a commercial breeder |
| Eurasian siskin | 2 | Trapped in Novosibirsk |

**Supplementary Table 2.** Size of expanded centromeres.

| Species | SC rank | SC length, $\mu\text{m}$ | Centromere length, $\mu\text{m}$ | Centromere length, % |
| --- | --- | --- | --- | --- |
| Flycatcher | SC1 | 26.2±1.9 | 2.3±0.4 | 8.6 |
| | SC2 | 20.3±2.3 | 2.2±0.7 | 11.2 |
| | SC3 | 21.9±3.9 | 2.5±0.4 | 11.6 |
| | SC5 | 15.0±1.9 | 3.9±0.7 | 25.9 |
| | SC6/7 | 13.4±1.6 | 2.1±0.6 | 15.4 |
| | SC8/10 | 8.8±1.5 | 2.3±0.4 | 26.3 |
| | SC11/13 | 6.8±0.4 | 2.8±0.3 | 41.3 |
| Canary | SC1 | 34.7±5.2 | 3.5±0.8 | 10.1 |
| | SC2/3 | 27.3±5.7 | 3.2±0.8 | 11.7 |
| | SC4 | 24.0±3.7 | 2.3±0.4 | 9.4 |
| | SC5 | 19.1±4.6 | 3.0±0.8 | 15.6 |
| | SC6 | 17.1±5.5 | 1.4±0.3 | 8.2 |
| | SC10/11/12 | 9.5±1.8 | 1.7±0.3 | 17.8 |
| Gouldian finch | SC1 | 27.9±4.4 | 3.6±0.6 | 13.6 |
| | SC2 | 22.5±3.6 | 4.4±0.6 | 19.8 |
| | SC3 | 22.3±3.7 | 3.8±0.7 | 17.1 |

## Notes

https://meiosislab.com/projects/chromosomes/17caryo.xls.

### Competing Interest Statement

The authors have declared no competing interest.

### Summary of Updates

One paragraph was added on the page 6, lines 143-146; Figure 3 revised.

